# Full-length Genome of a *Ogataea polymorpha* strain CBS4732 *ura3*Δ reveals large duplicated segments in subtelomeric regions

**DOI:** 10.1101/2021.04.17.440260

**Authors:** Jia Chang, Jinlong Bei, Qi Shao, Hemu Wang, Huan Fan, Tung On Yau, Wenjun Bu, Jishou Ruan, Dongsheng Wei, Shan Gao

**Affiliations:** Key Laboratory of Molecular Microbiology and Technology, Ministry of Education, College of Life Science, Nankai University, Tianjin, 300071, P. R. China; Guangdong Provincial Key Laboratory for Crop Germplasm Resources Preservation and Utilization, Agro-Biological Gene Research Center, Guangdong Academy of Agricultural Sciences, Guangzhou, Guangdong 510640, P. R. China; Guangdong Laboratory for Lingnan Modern Agriculture, Guangzhou, Guangdong 510642, P. R. China; Tianjin Hemu Health Biotechnological Co., Ltd, Tianjin, Tianjin 300384, P.R.China; John Van Geest Cancer Research Centre, School of Science and Technology, Nottingham Trent University, Nottingham, NG11 8NS, United Kingdom; Tianjin Institute of Animal Husbandry and Veterinary Research, Tianjin 300192, P.R.China; School of Mathematical Sciences, Nankai University, Tianjin, Tianjin 300071, P.R.China

**Keywords:** Methylotrophic yeast, Ogataea, DL-1, NCYC495, rDNA quadruple

## Abstract

**Background:** Currently, methylotrophic yeasts (e.g., *Pichia pastoris, Ogataea polymorpha*, and *Candida boindii*) are subjects of intense genomics studies in basic research and industrial applications. In the genus *Ogataea*, most research is focused on three basic *O. polymorpha* strains—CBS4732, NCYC495, and DL-1. However, the relationship between CBS4732, NCYC495, and DL-1 remains unclear, as the genomic differences between them have not be exactly determined without their high-quality complete genomes. As a nutritionally deficient mutant derived from CBS4732, the *O. polymorpha* strain CBS4732 *ura3*Δ (named HU-11) is being used for high-yield production of several important proteins or peptides. HU-11 has the same reference genome as CBS4732 (noted as HU-11/CBS4732), because the only genomic difference between them is a 5-bp insertion.

**Results:** In the present study, we have assembled the full-length genome of *O. polymorpha* HU-11/CBS4732 using high-depth PacBio and Illumina data. Long terminal repeat (LTR) retrotransposons, rDNA, 5’ and 3’ telomeric, subtelomeric, low complexity and other repeat regions were curated to improve the genome quality. Particularly, we detected large duplicated segments (LDSs) in the subtelomeric regions and exactly determined all the structural variations (SVs) between CBS4732 and NCYC495.

**New findings mainly include:** (1) the genomic differences between HU-11/CBS4732 and NCYC495 include single nucleotide polymorphisms, small insertions and deletions, and only three SVs; (2) six genes were incorporated into CBS4732 from *Cyberlindnera jadinii* by horizontal gene transfer and may bring HU-11/CBS4732 new biological functions or physiological properties; (3) many recombination events may have occurred on chromosome 4 and 5 of CBS4732 and NCYC495’ ancestors and two large segments were acquired by CBS4732 and NCYC495 from chromosome 6 and *C. jadinii* during recombination, respectively; and (4) the genome expansion in methylotrophic yeasts is mainly driven by large segment duplication in subtelomeric regions.

**Conclusions:** The present study preliminarily revealed the complex relationship between CBS4732, NCYC495, and DL-1. The new findings provide new opportunities for in-depth understanding of genome evolution in methylotrophic yeasts and lay the foundations for the industrial applications of *O. polymorpha* CBS4732, NCYC495, DL-1, and their derivative strains. The full-length genome of the *O. polymorpha* strain HU-11/CBS4732 should be included into the NCBI RefSeq database for future studies of *Ogataea* spp..

## Introduction

Currently, methylotrophic yeasts (e.g., *Pichia pastoris, Hansenula polymorpha*, and *Candida boindii*) are subjects of intense genomics studies in basic research and industrial applications. However, genomic research on *Ogataea (Hansenula) polymorpha* trails behind that on *P. pastoris* [1], although they both are popular and widely used species of methylotrophic yeasts. In the genus *Ogataea*, most research is focused on three basic *O. polymorpha* strains—CBS4732 (synonymous to NRRL-Y-5445 or ATCC34438), NCYC495 (synonymous to NRRL-Y-1798, ATCC14754, or CBS1976), and DL-1 (synonymous to NRRL-Y-7560 or ATCC26012). These three strains are of independent geographic and ecological origins: CBS4732 was originally isolated from soil irrigated with waste water from a distillery in Pernambuco, Brazil in 1959 [2]; NCYC495 is identical to a strain first isolated from spoiled concentrated orange juice in Florida and initially designated as *Hansenula angusta* by Wickerham in 1951 [3]; and DL-1 was isolated from soil by Levine and Cooney in 1973 [4]. CBS4732 and its derivatives—LR9, and RB11—have been developed as genetically engineered strains to produce many heterologous proteins, including enzymes (e.g., feed additive phytase), anticoagulants (e.g., hirudin and saratin), and an efficient vaccine against hepatitis B infection [5]. As a nutritionally deficient mutant derived from CBS4732 (CBS4732 *ura3*Δ), the *O. polymorpha* strain HU-11 [6] is being used for high-yield production of several important proteins or peptides, particularly including recombinant hepatitis B surface antigen (HBsAg) vaccine [7] and hirudin [8]. HU-11 has the same reference genome as CBS4732 (noted as HU-11/CBS4732), as the only genomic difference between them is a 5-bp insertion caused by frame-shift mutation of its *URA3* gene, which encodes orotidine 5’-phosphate decarboxylase. Although CBS4732 and NCYC495 are classified as *O. polymorpha*, and DL-1 is reclassified as *O. parapolymorpha* [9], the relationship between CBS4732, NCYC495, and DL-1 remains unclear, as the genomic differences between them have not be exactly determined due to lack of their high-quality complete genomes. Thus, the knowledge obtained from any of three strains can not be used to other strains.

To facilitate genomic research of yeasts, genome sequences have been increasingly submitted to the Genome-NCBI datasets. Among the genomes of 34 species in the *Ogataea* or *Candida* genus (**Supplementary file 1**), those of NCYC495 and DL-1 have been assembled at chromosome level. However, the other genomes have been assembled at the contig or scaffold level. Furthermore, the genome sequence of CBS4732 was not available in the Genome-NCBI datasets until this manuscript was drafted. Among the genomes of 33 *Komagataella (Pichia*) spp., the genome of the *P. pastoris* strain GS115 is the only genome assembled at chromosome level. The main problem of these *Ogataea, Candida*, or *Pichia* genomes is their incomplete sequences and poor annotations. For example, the rDNA sequence (GenBank: FN392325) of *P. pastoris* GS115 cannot be well aligned to its genome (Genbank assembly: GCA_001708105). Most genome sequences do not contain complete subtelomeric regions and, as a result, subtelomeres are often overlooked in comparative genomics [10]. For example, the genome of DL-1 has been analyzed for better understanding the phylogenetics and molecular basis of *O. polymorpha* [1]; however, it does not contain complete subtelomeric regions due to assembly using short sequences. Another problem of current yeast genome data is that the complete sequences of mitochondrial genomes is not simultaneously released with those of nuclear genomes. The only complete mitochondrial genome in the NCBI GenBank database is the *O. polymorpha* DL-1 mitochondrial genome (RefSeq: NC_014805). More high-quality complete genome sequences of *Ogataea* spp. need to be sequenced to bridge the gap in *Ogataea* basic research and industrial applications.

In the present study, we have assembled the full-length genome of *O. polymorpha* HU-11/CBS4732 using high-depth PacBio and Illumina data and conducted the annotation and analysis to achieve the following research goals: (1) to provide a high-quality and well-curated reference genome for future studies of *Ogataea* spp.; (2) to determine the relationship between CBS4732, NCYC495, and DL-1; and (3) to discover important genomic features (e.g., high yield) of *Ogataea* spp. for basic research (e.g., synthetic biology) and industrial applications.

## Results and Discussion

### Genome sequencing, assembly and annotation

One 500 bp and one 10 Kbp DNA library were prepared using fresh cells of *O. polymorpha* HU-11 and sequenced on the Illumina HiSeq X Ten and PacBio Sequel platforms, respectively, for *de novo* assembly of a high-quality genome. Firstly, 18,319,084,791 bp cleaned PacBio DNA-seq data were used to assembled the complete genome, except for the rDNA region (**analyzed in further detail in subsequent sections**), with an extremely high depth of ~1800X. However, the assembled genome using high-depth PacBio data still contained two types of errors in the low complexity (**Figure 1A**) and the short tandem repeat (STR) regions, respectively (**Figure 1B**). Then, 6,628,480,424 bp cleaned Illumina DNA-seq data were used to polish the complete genome of HU-11/CBS4732 to remove the two types of errors. However, Illumina DNA-seq data contained errors in the long (>10 copy numbers) poly(GC) regions. Following this, the poly(GC) regions, polished using Illumina DNA-seq data, were curated using PacBio subreads (**Figure 1C**). Finally, Long Terminal Repeat retrotransposons (LTR-rts), rDNA (**analysed in more details in following sections**), 5’ and 3’ telomeric, subtelomeric, low complexity, and other repeat regions were curated to obtain the full-length genome using 103,345 long (> 20 Kbp) PacBio subreads (**Supplementary file 1**).

**Figure 1.**
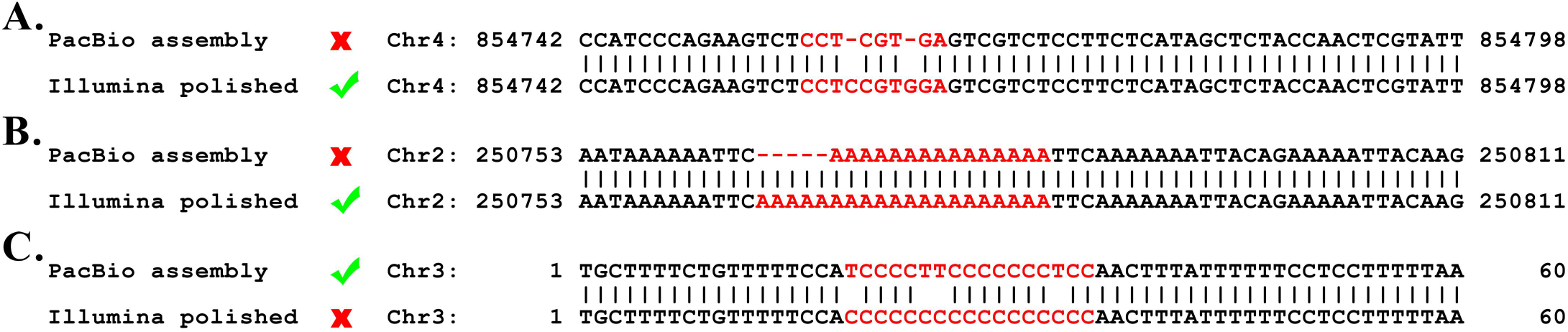
Errors in PacBio data and Illumina data. The errors in the low complexity and short tandem repeat (STR) regions can be corrected during the genome polishment using Illumina data, while the errors in the long (>10 copy numbers) poly(GC) regions need be curated using PacBio data after the genome polishment. **A.** A example to show that the assembled genomes using high-depth PacBio data still contain errors in the low complexity regions. **B.** A example to show that the assembled genomes using high-depth PacBio data still contain errors in the STR regions. **C.** A example to show that the genome polishment using Illumina data causes errors in the long poly(GC) regions.

*Ogataea polymorpha* HU-11/CBS4732 has a nuclear genome (**Figure 2A**) with a summed sequence length of 9.1 Mbp and a mitochondrial (mt) genome (**Figure 2B**) with a sequence length of 59,496 bp (**Table 1**). For the data submission to the GenBank database, the sequence of circular mt genome was anticlockwise linearized, starting at the first nt of large subunit ribosomal RNA (rrnL). Analysis of long PacBio subreads revealed that the telomeric regions at 5’ and 3’ ends of each chromosome consist of tandem repeats (TRs) [ACCCCGCC]_n_ and [GGCGGGGT]_n_ (n is the copy number) with average lengths of 166 bp and 168 bp (~20 copy numbers), respectively. As these TRs vary in lengths, the 5’ and 3’ telomeric regions were not included into the seven linear chromosomes of HU-11/CBS4732, which were named as 1 to 7 from the smallest to the largest, respectively (**Table 1**). The full-length *O. polymorpha* HU-11/CBS4732 genome includes the complete sequences of all seven chromosomes, while the 5’ and 3’ ends of NCYC495 (**RefSeq: NW_017264698-704**) or DL-1 chromosomes (**RefSeq: NC_027860-66**) have many errors (**Supplementary file 1**). Recently, a new project has been conducted to provide a high-quality reference genome of DL-1 (**GenBank: CP080316-22**) based on Nanopore technology. Therefore, we recommend the inclusion of our genome sequences into the NCBI RefSeq database to facilitate future studies on *O. polymorpha* CBS4732 and its derivatives-LR9, RB11, and HU-11.

**Figure 2.**
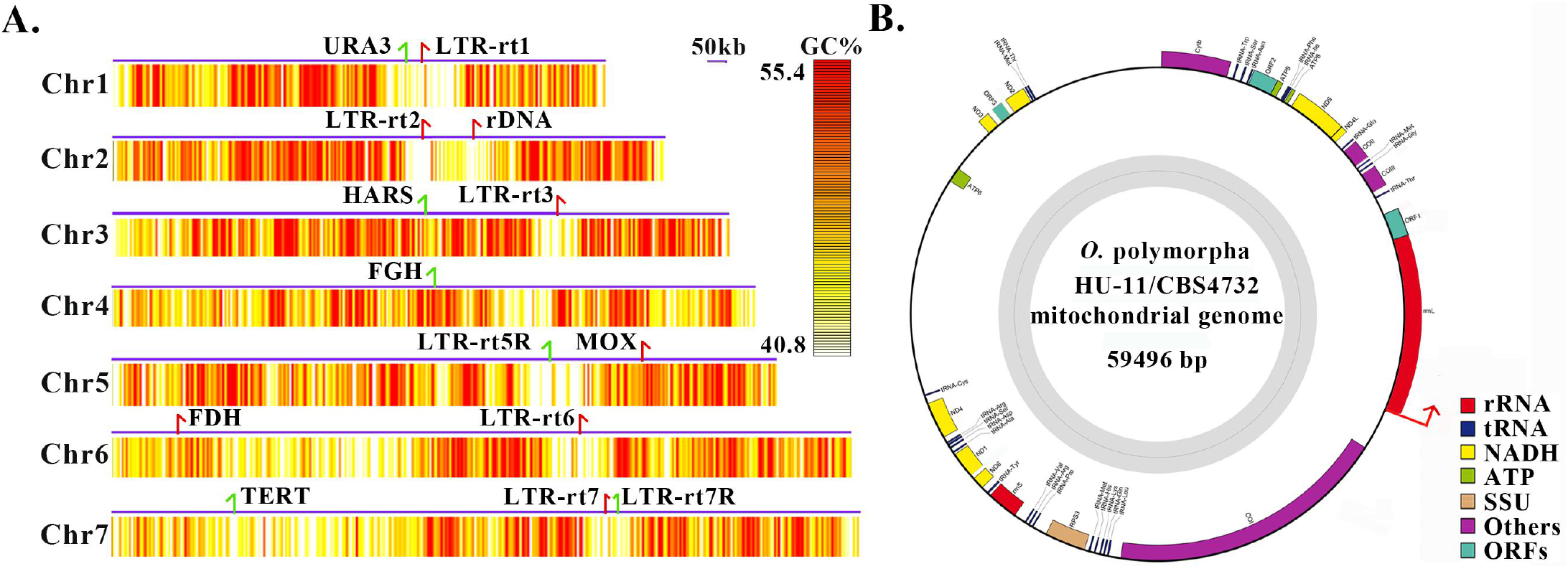
Full-length genome of the *Ogataea polymorpha* strain HU-11. **A.** The full-length *O. polymorpha* HU-11/CBS4732 genome includes the complete sequences of seven linear chromosomes, which were named as 1 to 7 from the smallest to the largest. The 5’ and 3’ telomeric regions were not included. The minimum, Q90, Q75, Q50, Q25, Q10 and maximum of GC contents (%) are 0.08, 0.408 0.436, 0.472, 0.514, 0.554 and 0.732. The GC contents (%) were calculated by 500-bp sliding windows and then trimmed between Q10 and Q90 for plotting the heatmaps. Long terminal repeat retrotransposons (LTR-rts) are indicated by red arrows (red and green colours represent sense and antisense strands) in the chromosomes. Markers genes indicated by red arrows (red and green colours represent sense and antisense strands) include URA3 (encoding orotidine 5'-phosphate decarboxylase), HARS (Hansenula autonomously replicating sequence), FGH (S-formylglutathione hydrolase), MOX (methanol oxidase), FDH (Formate dehydrogenase) and TERT (telomerase reverse transcriptase). **B.** For the data submission to the GenBank database, the genome sequence of circular mitochondrion was anticlockwise linearized, starting at the first nt (indicated by a red arrow) of rrnL, which may include a part of the control region. SSU: small subunit; RPS3: ribosomal protein S3; rrnL: large subunit ribosomal RNA; rrnS: small subunit ribosomal RNA.

**Table 1.** Genomes of three basic *O. polymorpha* strains.

| Chromosome | CBS4732/HU-11 | NCYC495 | DL-1 | HU-11 Size (bp) | Marker |
| --- | --- | --- | --- | --- | --- |
| Chr1 | CP073033 | NW_017264703 | NC_027865 | 1,000,895 | URA3 |
| Chr2 | CP073034 | NW_017264704 | NC_027866 | 1,125,341 | rDNA |
| Chr3 | CP073035 | NW_017264702 | NC_027864 | 1,265,401 | HARS |
| Chr4 | CP073036 | NW_017264701 | NC_027863 | 1,315,956 | FGH |
| Chr5 | CP073037 | NW_017264700 | NC_027862 | 1,357,435 | MOX |
| Chr6 | CP073038 | NW_017264698 | NC_027860 | 1,513,391 | FDH |
| Chr7 | CP073039 | NW_017264699 | NC_027861 | 1,525,912 | TERT |
| ChrM | CP073040 | NA | NC_014805 | 59,496 | COIII |
| Total (Mbp) | 9.1 | 8.97 | 8.87 |  |  |
| GC% | 47.76 | 47.86 | 47.83 |  |  |
| Gene/mRNA# | 5138* | 5138* | 5325* |  |  |
| tRNA# | 80 | 80 | 80 |  |  |
| rRNA# | 4×20 | 4×6 | 4×25 |  |  |

The 9.1 Mbp length of the HU-11/CBS4732 genome is close to the estimated length of the *O. polymorpha* DL-1 genome [1], while the published NCYC495 and DL-1 genomes (**Table 1**) have shorter lengths of 8.97 and 8.87 Mbp, respectively (**Table 1**), thus they need be further completed. The GC contents of the HU-11, NCYC495, and DL-1 genomes are comparable (~48%). Taking advantage of the full-length HU-11/CBS4732 genome sequence for the genome annotation and comparison, we determined the exact location of the rDNA genes and LTR-rts in seven chromosomes (**Figure 2A**) and detected large duplicated segments (LDSs) in the subtelomeric regions (**described in more detail in succeeding sections**). Syntenic comparison (**Methods and Materials**) revealed that *O. polymorpha* NCYC495 is so phylogenetically close to HU-11/CBS4732 that the syntenic regions covers nearly 100% of their genomes, whereas DL-1 is significantly distinct from HU-11/CBS4732. Using syntenic regions in the full-length HU-11/CBS4732 genome, we corrected the genome sequences of NCYC495 (**RefSeq: NW_017264698-704**) and DL-1 (**RefSeq: NC_027860-66**). Using a high quality RNA-seq data of NCYC495 (NCBI SRA: SRP124832), we improved the gene annotations of HU-11/CBS4732, NCYC495, and DL-1 (**Table 1**): (1) HU-11/CBS4732 has 5,138 protein-coding genes, including 4,716 single exon genes, and 422 multiple exon genes; (2) NCYC495 has 5,138 protein-coding genes, including 4,714 single exon genes, and 424 multiple exon genes; (3) DL-1 has 5,325 protein-coding genes, including 4,861 single exon genes, and 464 multiple exon genes; and (4) HU-11/CBS4732, NCYC495, and DL-1 have 80 identical tRNA genes.

### Organization of rDNA genes

An rDNA TR of HU-11/CBS4732, NCYC495, or DL-1 encodes 5S, 18S, 5.8S, and 25S rRNAs (named as quadruple in the present study), with a length of ~8,100 bp (**Supplementary file 1**). The copy number of rDNA TRs was estimated as 20 in the HU-11/CBS4732 genome (**Figure 3A**), while that was estimated as 6 and 25 in NCYC495 and DL-1, respectively [1]. TRs of HU-11/CBS4732 and NCYC495 rDNAs share a very high nucleotide (nt) sequence identity of 99.5% (8,115/8,152), while those of HU-11/CBS4732 and DL-1 rDNAs share a comparatively low nt sequence identity of 97% (7,530/7,765). As the largest TR region (~162 Kbp) in the HU-11/CBS4732 genome, the only rDNA locus is located on chromosome 2 and the organization of rDNA genes with different copy numbers may be conserved in the *Ogataea* genus. A rDNA TR of *Saccharomyces cerevisiae* also contains 5S, 18S, 5.8S, and 25S rDNAs as a quadruple, repeating two times on chromosome 7 of its genome (**Figure 3B**). Four other 5S rDNAs are located separately away from the rDNA quadruples in *S. cerevisiae*. Compared to *O. polymorpha* or *S. cerevisiae* with only one rDNA locus, *Pichia pastoris* GS115 carries several rDNA loci, which are interspersed in three of its four chromosomes. Since the genome of *P. pastoris* GS115 (Genbank assembly: GCA_001708105) is incomplete and poorly annotated, we estimated the copy number of its rDNAs as three. In eukaryotes, rDNAs encoding 18S, 5.8S, and 28S rRNAs that are transcribed into a single RNA precursor by RNA polymerase I are also organized in TRs. For example, there are approximately 200–600 rDNA copies (**Figure 3C**) distributed in short arms of the five acrocentric chromosomes (chromosomes 13, 14, 15, 21, and 22) of human. [11]. In prokaryotic cells, 5S, 23S, and 16S rRNA genes are typically organized as a co-transcribed operon. There may be one or more copies of the operon dispersed in the genome and the copy numbers typically range from 1 to 15 in bacteria. For example, there are four copies at two rDNA loci in chromosome 1 (GenBank: CP022603) and 2 (GenBank: CP022604) of *Ochrobactrum quorumnocens* (**Figure 3D**). Compared to those of *S. cerevisiae*, human, and bacteria rDNAs (**Figure 3BCD**), 20 copies of *O. polymorpha* rDNA quadruples are very closely organized, suggesting that their transcription is regulated with high efficiency. This genomic feature may contribute to the high yield characteristics of *O. polymorpha.*

**Figure 3.**
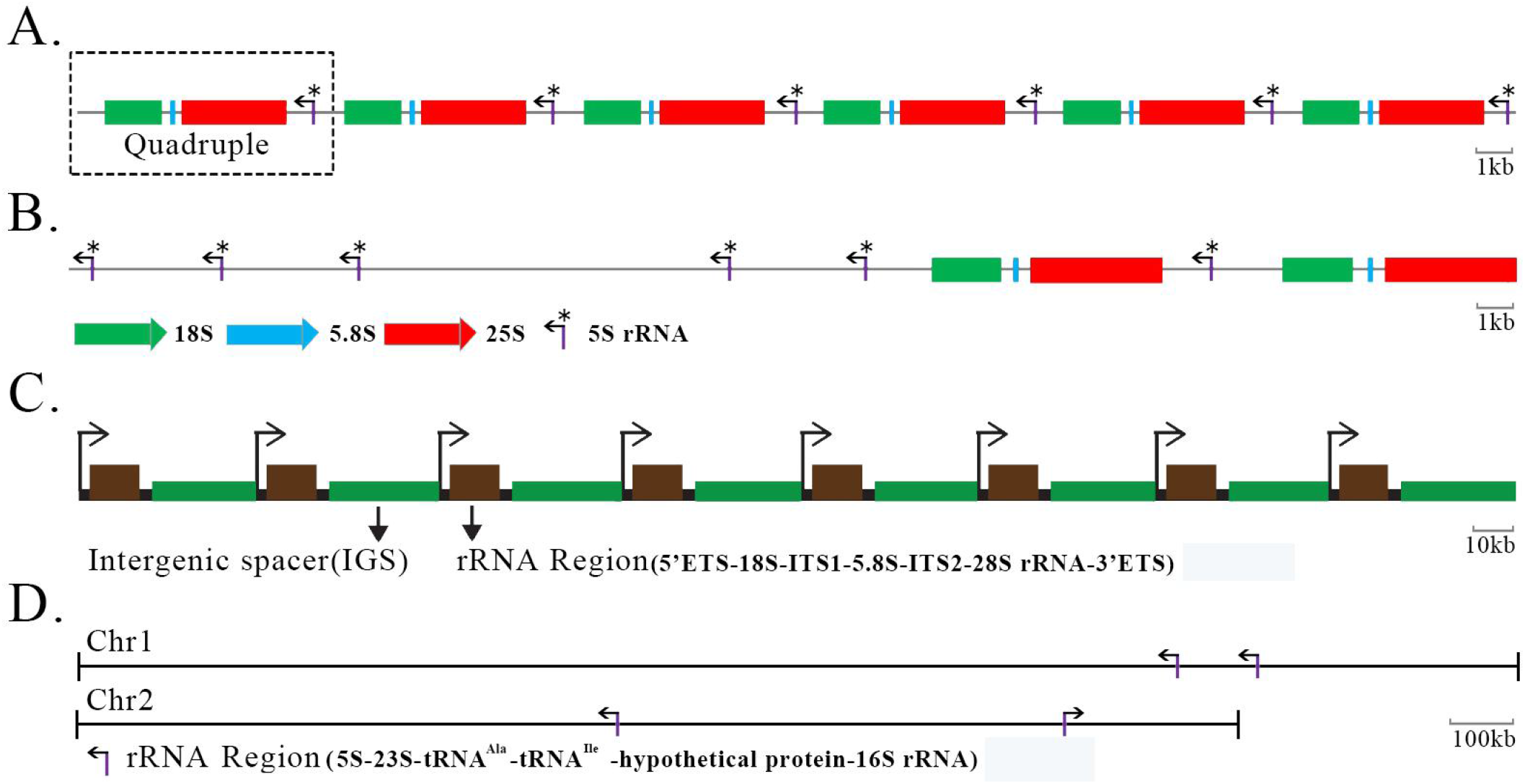
Organization of rDNA genes in yeasts, human and bacteria. **A.** The only rDNA locus is located in chromosome 2 (GenBank: CP073034) of the Ogataea polymorpha strain HU-11, containing 20 copies of TRs. Here only six copies of TRs are shown. B. An rDNA TR of Saccharomyces cerevisiae also contains 5S, 18S, 5.8S and 25S rDNAs as a quadruple, repeating 2 times on chromosome 7 of its genome. Four other 5S rDNAs are located separately away from the rDNA quadruples in S. cerevisiae. C. Each human rDNA unit has an rRNA region and an intergenic spacer (IGS). Here only eight units are shown. ITS: internal transcribed spacer; ETS: external transcribed spacers. D. There are four copies of rRNA regions at two rDNA loci on chromosome 1 (GenBank: CP022603) and 2 (GenBank: CP022604) of the Ochrobactrum quorumnocens genome.

Besides the high similarity of genomic arrangement, the rDNAs of *S*. *cerevisiae* and *O. polymorpha* HU-11/CBS4732 share high nt sequence identities of 95.3% (1720/1805), 96.2% (152/158), 92% (3,111/3,381), and 96.7% (117/121) for 18S, 5.8S, 25S, and 5S rDNAs, respectively. However, the rDNAs (Genbank: FN392325) of *P. pastoris* GS115 and *O. polymorpha* HU-11/CBS4732 have nt sequence identities of 87.3% (1477/1691), 80% (84/105), and 80.5% (2,073/2,576) for 18S, 5.8S, and 25S rDNAs, respectively. This finding contradicts the results of a previous study [1] in which phylogenetic analysis using 153 protein-coding genes showed that *Pichia pastoris* GS115 and *O. polymorpha* are members of a clade that is distinct from the one that *S. cerevisiae* belongs to. However, the present study revealed that HU-11/CBS4732 is phylogenetically closest to NCYC495, followed by DL-1, *S. cerevisiae*, and *P. pastoris* GS115, if we use rDNAs for the phylogenetic analysis. In addition, the present study showed that the rDNA genes are more conservative than the protein-coding genes in yeasts, and rDNA is an important feature of yeasts for their detection, identification, classification and phylogenetic analysis.

### Long terminal repeat retrotransposons

LTRs with lengths of 322 bp were discovered in all seven chromosomes of HU-11/CBS4732. These LTRs with low GC content of 29% (94/322) are flanked by TCTTG and CAACA at their 5’ and 3’ ends (**Figure 4A**). All the LTRs in HU-11/CBS4732 were identified as components of Tpa5 LTR-rts (GenBank: AJ439553) from *Pichia angusta* CBS4732 (a former name of *O. polymorpha* CBS4732) in a previous study. A LTR-rt consists of 5’ LTR, 3’ LTR, and a single open reading frame (ORF) encoding a putative polyprotein (**Figure 4A**). This polyprotein, if translated, can be processed into truncated Gag (GAG), protease (PR), integrase (IN), reverse transcriptase (RT), and RNase H (RH). Based on the gene order (PR, IN, RT, and RH), the LTR-rts of HU-11/CBS4732 were classified into the Ty5 type of the Ty1/copia group (Ty1, 2, 4, and 5 types) [12].

**Figure 4.**
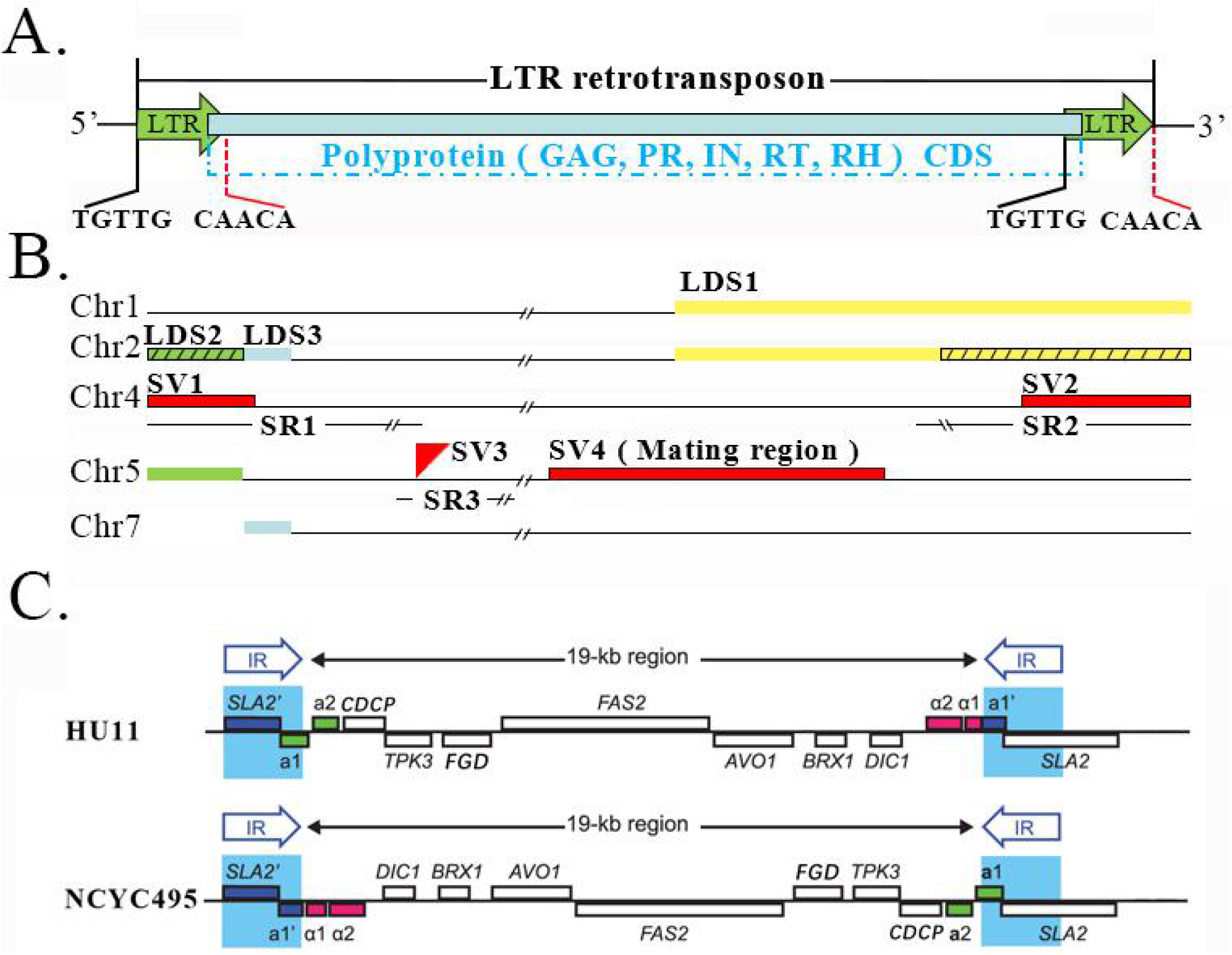
LTR retrotransposons, large duplicated segments and structural variations. **A.** NCYC495 and HU-11/CBS4732 share identical 322-bp LTRs, which are flanked by TCTTG and CAACA at their 5’ and 3’ ends. Three of seven LTR-rts of HU-11/CBS4732 does not have homologs in the NCYC495 genome due to misassembly. A LTR-rt consists of 5’ LTR, 3’ LTR and a single open reading frame (ORF) encoding a putative polyprotein. This polyprotein, if translated, can be processed into trunculated gag (GAG), protease (PR), integrase (IN), reverse transcriptase (RT) and RNase H (RH). **B.** Chr1, 2, 4, 5 and 7 represent the chromosomes (GenBank: CP073033, 34, 36, 37 and 39) in the HU-11/CBS4732 genome. Three large duplicated segments (LDSs) named LDS1 (in yellow color), 2 (in green color) and 3 (in blue color) are supposed to be included in both NCYC495 and HU-11/CBS4732 genomes. However, LDS2 and a 14,090 bp part of LDS1's homolog (indicated by black slashs) were not assembled into chromosome 2 in the NCYC495 genome. The genomic differences between the HU-11/CBS4732 and NCYC495 only include three structural variations (SVs), named SV1, 2 and 3 (in red color). The three SVs are located in three large syntenic regions (SRs) of HU-11/CBS4732, NCYC495, and DL-1 genomes with very high nt sequence identities, named SR1, 2 and 3. **C.** The graphic elements used to represent the genomes and genes were originally used in the previous study [9]. SV4 is a 22.6-Kb DNA region which functions in the determination of the yeast mating-type (MAT). The HU-11/CBS4732 genome (GenBank: CP073033-40) contains a 22.6-Kb MAT region where MATα can be transcribed, while the NCYC495 genome (RefSeq: NW_017264698-704) contains an identical 22.6-Kb MAT region where MATa can be transcribed.

With the length corrected from 4,883 bp to 4,882 bp, a sequence (GenBank: AJ439558) was used as reference of Tpa5 LTR-rts to search for homologs. The results confirmed that HU-11/CBS4732 is phylogenetically closest to NCYC495 and they share identical 322-bp LTRs. However, the 322-bp LTRs of HU-11/CBS4732 and NCYC495 are quite distinct from the 282-bp LTRs of DL-1, which were reported as 290-bp solo LTRs in the previous study [1]. In addition, the amino acid (aa) sequences of the polyprotein with the length of 1417-aa in HU-11/CBS4732 and NCYC495 LTR-rts are distinct from those in DL-1. Based on the records in the UniProt Knowledgebase (UniProtKB), *O. polymorpha* strains DL-1, ATCC26012, BCRC20466, JCM22074, and NRRL-Y-7560 have nearly the same aa sequences (**UniProt: W1QI12**) of the polyprotein. These results suggest that the LTR-rt is another important feature of yeasts useful for their detection, identification, classification, and phylogenetic analysis. Using RNA-seq data of NCYC495 (SRA: SRP124832), we discovered that the polyproteins in the LTR-rts of *O. polymorpha* are transcribed. If these polyproteins can be translated for biological functions merits further studies.

In the previous study, 50,000 fragments of 13 *Hemiascomycetes* species were used to identify LTR-rts. However, the analysis was probably biased as it was based on only random sequences of approximately 1 kb on an average and not the complete genome sequences [12]. In the present study, seven copies of intact LTR-rts (**Supplementary file 1**) were discovered and accurately positioned in the HU-11/CBS4732 genome (**Figure 2A**); five of them are located on the sense strands of chromosome 1, 2, 3, 6, and 7 (named LTR-rt1, 2, 3, 6, and 7), while the other two are located on the antisense strands of chromosome 5 and 7 (named LTR-rt5R and 7R). LTR-rt1, 3, and 6 share very high nt identities of 99.9% with each other. LTR-rt1 or 3 contains a single ORF encoding a polyprotein with the same aa sequence, while LTR-rt6 contains a single ORF with a 42-bp insertion (encoding RSSLFDVPCSPTVD), compared to LTR-rt1 and 3. LTR-rt2, 5R, 7, and 7R contain several single nucleotide polymorphisms (SNPs), small insertions and deletions (InDels), which break the single ORFs into several ORFs. The homologs of LTR-rt2, 3, and 5R in HU-11/CBS4732 are present in the NCYC495 genome with very high nt identities of 99.9%. The homologs of LTR-rt1, 7, and 7R, however, were not detected in the NCYC495 genome. Further analysis determined that their absence in the NCYC495 genome was resultant from misassembly (**described in more detail subsequently**).

### Structural variation and large segment duplication

Sequence comparison between the NCYC495 and HU-11/CBS4732 genomes showed that they share a nt identity of 99.5% through the whole genomes, including the rDNA region and LTR-rts. However, the DL-1 and HU-11/CBS4732 genomes share a comparatively low nt identity (< 95%) through the whole genomes. Syntenic comparison revealed that NCYC495 is so phylogenetically similar to HU-11/CBS4732 that the syntenic regions cover nearly 100% of their genomes, whereas DL-1 is significantly distinct from HU-11/CBS4732 (**As shown in preceding sections**). Subsequently, the detection of structural variations (SVs) was performed between the NCYC495 and HU-11/CBS4732 genomes. Further analysis revealed that all detected SVs are large InDels (two types of SVs) and most of them are errors in the assembly of NCYC495 genome (**Figure 4B**), particularly including: (1) LTR-rt1, 7, and 7R (absent in NCYC495) need be included in the NCYC495 genome; (2) two large deletions (absent in NCYC495) need be added at 5’ and 3’ ends of chromosome 2; and (3) a large insertion (absent in HU-11/CBS4732) is an over-assembled segment at 3’ end of chromosome 6 (**NW_017264698:1509870-1541475**), which need be removed from chromosome 6. Telomeric TRs [GGCGGGGT]_n_(NW_017264698:1509840-1509869) were discovered at 5’ end of this over-assembled segment, confirming that it resulted from misassembly.

The main reason to cause the above assembly errors in the NCYC495 genome is the misassembly of LDSs in the subtelomeric regions and LTR-rts, which resulted in false-positive SVs. These LDSs and LTR-rts (**described above**) were correctly assembled in the HU-11/CBS4732 genome. Using long (> 30 Kb) PacBio subreads, human curation was performed to verify the locations of the LDSs, particularly three LDSs named LDS1, 2 and 3 (**Figure 4B**). LDS1 and its homolog are present at 3’ ends of chromosome 1 and 2 in the HU-11/CBS4732 genome, respectively. There are only four mismatches and one 1-bp gap between LDS1 and its homolog with a length of 27,850 bp. In the NCYC495 genome, LDS1 was correctly assembled into 3’ end of chromosome 1, but a 14,090 bp part of LDS1's homolog was not assembled into 3’ end of chromosome 2, which corresponds to a large deletion (**described above**). LDS2 and its homolog are present at 5’ ends of both chromosomes 2 and 5 in the HU-11/CBS4732 genome with a length of approximate 5,100 bp, while the homolog of LDS2 was correctly assembled into 5’ end of chromosome 5, but LDS2 was not assembled into 5’ end of chromosome 2 in the NCYC495 genome, which corresponds to the other large deletion (**described above**). LDS3 is downstream of LDS2 on chromosome 2 in the HU-11/CBS4732 genome with a length of approximate 2,500 bp, and the homolog of LDS3 is present at 5’ end of chromosome 7. Different from LDS1 and LDS2, LDS3 and its homolog were correctly assembled in the NCYC495 genome. As an important finding, telomeric TRs [ACCCCGCC]_n_ (n > 2) were discovered at 3’ ends of LDS2 and its homolog (located on both chromosomes 2 and 5), and at 3’ end of LDS3's homolog (located on chromosome 7); this finding indicated that 3’ ends of these LDSs were integrated at 5’ ends of telomeric TRs.

### Genomic differences between HU-11/CBS4732 and NCYC495

After correction of all assembly errors in the NCYC495 genome, syntenic regions covered the whole HU-11/CBS4732 and NCYC495 genomes except four SVs, named SV1, SV2, SV3, and SV4 (**Figure 4B**). Further analysis confirmed that the genomic differences between HU-11/CBS4732 and NCYC495 include SNPs, small InDels, and only three SVs (SV1, SV2, and SV3), not SV4. SV4 is a 22.6-Kb DNA region (**Figure 4C**) which functions in the determination of the yeast mating-type (MAT). Yeast mating generally occurs between two haploid cells with opposite genotypes (MATa and MATα) at this locus, to form a diploid zygote (MATa/α). *O. polymorpha* chromosome 5 contains both a MATa locus and a MATα locus, approximately 19 Kb apart (**Figure 4C**). The two MAT loci are beside two copies of an identical 2-Kb DNA sequence, which form two inverted repeats (IRs). During MAT switching, the two copies of the IR recombine, inverting the orientation of the 19-Kb region relative to the rest of the chromosome. The MAT locus proximal to the centromere is not transcribed, probably due to silencing by centromeric heterochromatin, whereas the distal MAT locus is transcribed [9]. The HU-11 genome (**GenBank: CP073033-40**) contains a 22.6-Kb MAT region (MAT-HU11) where MATα can be transcribed, while the NCYC495 genome (**RefSeq: NW_017264698-704**) contains an identical 22.6-Kb MAT region (MAT-NCYC495) where MATa can be transcribed. There are only one 1-bp gap between the large segments MAT-HU11 and MAT-NCYC495 (**Supplementary file 1**). Using long PacBio subreads, we found that MAT switching rarely occurred in HU-11 under normal conditions. MAT regions can not be used as a genomic marker to characterize different *O. polymorpha* strains.

SV1 and SV2 are present at 5’ ends and 3’ ends of chromosome 4 in both HU-11/CBS4732 and NCYC495 genomes, respectively, while the location of SV3 is close to 5’ ends of chromosome 5 (**Figure 4C**). Five sequences involved in these three SVs are SV1-CBS4732 and SV2-CBS4732 in the HU-11/CBS4732 genome and NCYC495, SV2-NCYC495, and SV3-NCYC495 in the NCYC495 genome (**Supplementary file 1**). These five sequences can be used to identify *O. polymorpha* strains, particularly CBS4732, NCYC495, and their derivative strains. Blasting the five sequences to the NCBI NT database, we found that SV1-CBS4732 and SV2-NCYC495 are nearly identical (>98%) to their homologs at 5’ and 3’ ends of chromosome 4 in the DL-1 genome (**GenBank: CP080319**), respectively, while SV1-NCYC495 and SV2-CBS4732 have no homologs on chromosome 4. As an insertion into the NCYC495 genome, SV3-NCYC495 has a very high nt sequence identity (>91%) to its homolog in the DL-1 genome. Further analysis showed that the three SVs are located in three large syntenic regions (SRs) of HU-11/CBS4732, NCYC495, and DL-1 genomes with very high nt sequence identities (>95%). Three SRs are: (1) SR1 with a length of 161,844 bp at 5’ ends of chromosome 4; (2) SR2 with a length of 81,748 bp at 3’ ends of chromosome 4; and (3) SR3 with a length of 11,087 bp close to 5’ ends of chromosome 5. These findings revealed that many recombination events occurred on chromosome 4 of CBS4732 and NCYC495’ ancestors, particularly: (1) recombination events occurred at 5’ end of chromosome 4 of the NCYC495’ ancestor, resulting in the acquisition of SV1-NCYC495; (2) recombination events occurred at 3’ end of chromosome 4 of the CBS4732’ ancestor, resulting in the acquisition of SV2-CBS4732; (3) recombination events occurred close to 5’ end of chromosome 5 of the CBS4732’ ancestor, resulting in the loss of SV3-HU11 (the homolog of SV3-NCYC495).

Only a few genes (predicted as 25) were involved in the three SVs between HU-11/CBS4732 and NCYC495 (**Table 2**). Blasting the proteins encoded by these 25 genes (**Supplementary file 1**) to the UniProt database, we found that the proteins (OGAPODRAFT_24127, 16381, 24129, 12876, and 16382) encoded in SV1-NCYC495 have the highest sequence similarities to their homologs (HPODL_02403, 02402, 02403, 02404, 02405, and 02398) at 3’ end of chromosome 6 (**RefSeq: NC_027860**) in the DL-1 genome. The proteins encoded by six genes (OGAPOHU_00003-08) in a major part (more than 80%) of SV2-CBS4732 have the highest sequence similarities to their homologs in the genome of the *Cyberlindnera jadinii* strain CBS1600. These findings indicated that SV1-NCYC495 from chromosome 6 and SV2-CBS4732 from *Cyberlindnera jadinii* were acquired by chromosome 4 of NCYC495 and CBS4732 via recombination, respectively. Furthermore, we found that the proteins encoded by two genes (OGAPOHU_00001-02) in the minor part of SV2-CBS4732 (from chromosome 4) have the highest sequence similarities to their homologs on other chromosomes. This revealed more combination events occurred between chromosome 4 and other genomes. Among 25 different genes between HU-11/CBS4732 and NCYC495, 10 and 11 genes were lost by NCYC495 and HU-11/CBS4732 via recombination, respectively (**Table 2**) and two genes were significantly changed, resulting in different aa sequences. As an important finding, six genes (OGAPOHU_00003-08) encoded in SV2-CBS4732 (**Table 2**) were incorporated into CBS4732 from *C. jadinii* by horizontal gene transfer (HGT) and may bring CBS4732 new biological functions or physiological properties, which are different from those of NCYC495 or DL-1.

**Table 2.**
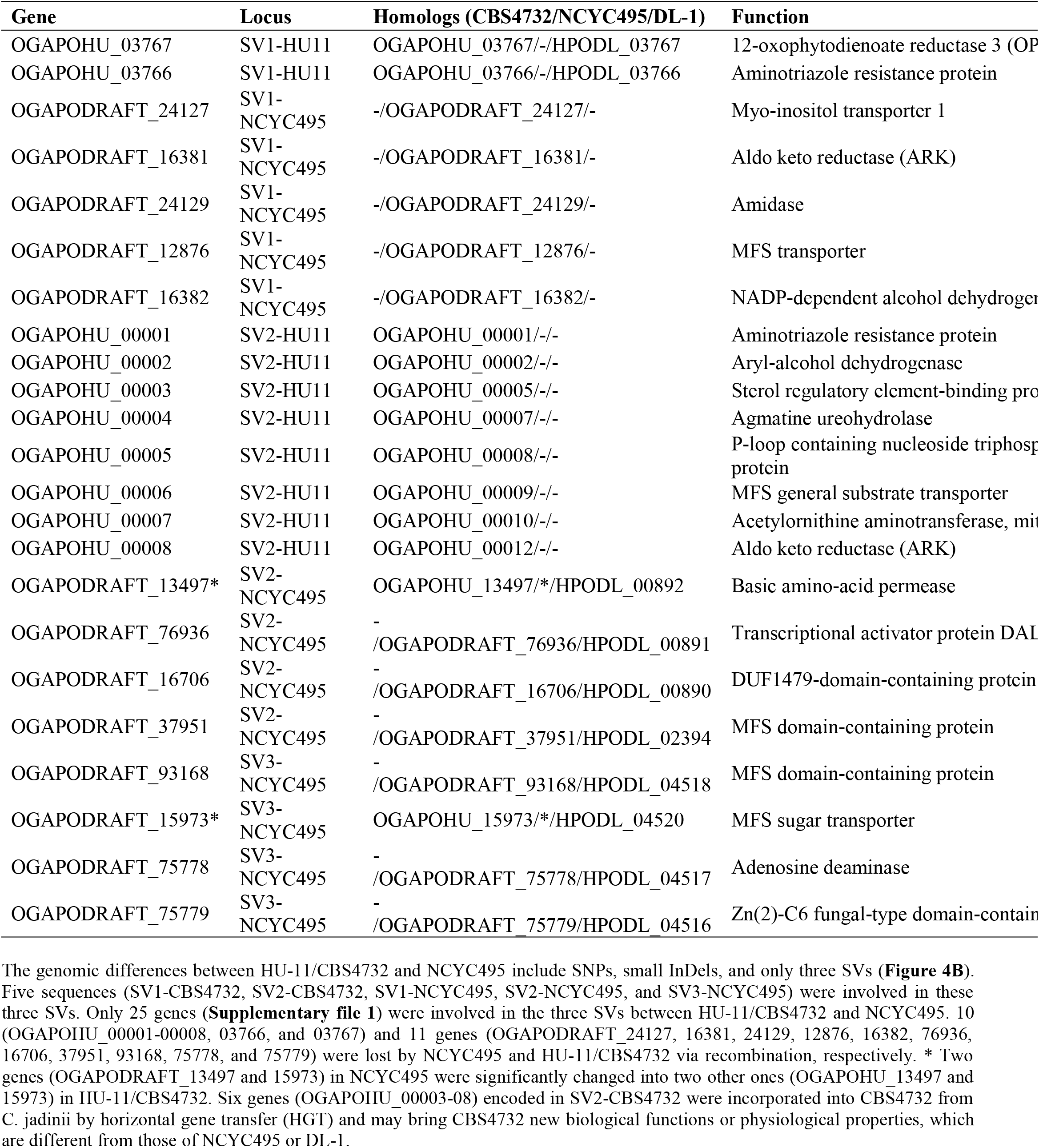
25 different genes between HU-11/CBS4732 and NCYC495.

## Conclusions

The *O. polymorpha* strain CBS4732 *ura3*Δ (named HU-11) is a nutritionally deficient mutant derived from CBS4732 by a 5 bp insertion of “GAAGT” into the 32^st^ position of the *URA3* CDS; this insertion causes a frame-shift mutation of the *URA3* CDS, resulting in the loss of the *URA3* functions. Since the difference between the genomes of CBS4732 and HU-11 is only five nts, HU-11 has the same reference genome as CBS4732 (HU-11/CBS4732). In the present study, we have assembled the full-length genome of *O. polymorpha* HU-11/CBS4732 using high-depth PacBio and Illumina data. Long Terminal Repeat retrotransposons (LTR-rts), rDNA, 5’ and 3’ telomeric, subtelomeric, low complexity, and other repeat regions were curated to improve the genome quality. Therefore, the full-length genome of the *O. polymorpha* strain HU-11/CBS4732 can be used as a reference for future studies of *Ogataea* spp.. For example, we corrected assembly errors in the NCYC495 genome using the full-length genome of HU-11/CBS4732, which facilitated in obtaining the full-length genome of NCYC495.

*O.* polymorpha NCYC495 is so phylogenetically close to HU-11/CBS4732 that the syntenic regions covers nearly 100% of their genomes. The genomic differences between HU-11/CBS4732 and NCYC495 include SNPs, small InDels, and only three SVs. Large segments SV1-CBS4732, SV2-CBS4732, SV1-NCYC495, SV2-NCYC495, and SV3-NCYC495 involved in the three SVs can be used to identify *O. polymorpha* strains, particularly CBS4732, NCYC495, and their derivative strains. As an important finding, six genes encoded in SV2-CBS4732 were incorporated into CBS4732 from *C. jadinii* by HGT and may bring HU-11/CBS4732 new biological functions or physiological properties, which are different from those of NCYC495 or DL-1. Many recombination events may have occurred on chromosome 4 and 5 of CBS4732 and NCYC495’ ancestors and two large segments (SV1-NCYC495 from chromosome 6 and SV2-CBS4732 from *C. jadinii*) were acquired by chromosome 4 of NCYC495 and CBS4732 via recombination, respectively. Recombination events occurred close to 5’ end of chromosome 5 of CBS4732’ ancestor, resulting in the loss of SV3-HU11 (the homolog of SV3-NCYC495) in the CBS4732 genome.

Using the high-quality full-length HU-11/CBS4732 genome, LDSs in subtelomeric regions were first discovered in methylotrophic yeast genomes, which was overlooked in the previous studies due to lack of PacBio or Nanopore sequencing. A computational study showed that subtelomeric families are evolving and expanding much faster than those which do not contain subtelomeric genes in yeasts. This study thus, indicated that the extraordinary instability of eukaryotic subtelomeres supports rapid adaptation to novel niches by promoting gene recombination and duplication followed by functional divergence of the alleles [10]. Our results suggest that the genome expansion in methylotrophic yeasts is mainly driven by large segment duplication in subtelomeric regions, accounting for the faster evolution and expension of subtelomeric gene families. The discovery of telomeric TRs at 3’ ends of these segments indicated that 3’ ends of these LDSs were integrated at 5’ ends of telomeric TRs. However, the underlying molecular mechanism (if via recombination or not) is still unknown.

## Methods and Materials

The *Ogataea polymorpha* strain HU-11 was obtained from Tianjin Hemu Health Biotechnological Co., Ltd. DNA extraction and quality control were performed as described in our previous study [13]. A 500 bp DNA library was constructed as described in our previous study [13] and sequenced on the Illumina HiSeq X Ten platform. A 10 Kb DNA library was constructed and sequenced on the PacBio Sequel platforms, according to the manufacturer's instruction. The software SMRTlink v5.0 (--minLength=50, -- minReadScore=0.8) was used for PacBio data cleaning and quality control, while the software Fastq_clean v2.0 [14] was used for Illumina data cleaning and quality control. The software MECAT v1.2 was used to assemble the HU-11/CBS4732 draft genome using PacBio data. To polish the HU-11/CBS4732 genome, Illumina data was aligned to the HU-11/CBS4732 draft genome using the software BWA. Then, the software samtools was used to obtain the BAM and pileup files from the alignment results. Perl scripts were used to extract the consensus sequence from the pileup file. This procedure was repeatedly performed to obtain the final genome sequence. The curation of genome and genes was performed using the software IGV. The software blast v2.9.0 was used to for syntenic comparison and SV detection. Statistical computation and plotting were performed using the software R v2.15.3 with the Bioconductor packages [15].

Syntenic comparison of genomes were performed using the CoGe website (https://genomevolution.org/CoGe). Among The genomes sequences of 34 species in the *Ogataea* or *Candida* genus were downloaded from the Genome-NCBI datasets and their accession numbers were included in **Supplementary file 1**. The reference genomes of *O. polymorpha* HU-11/CBS4732, NCYC495 and DL-1 are available at the NCBI GenBank or RefSeq database under the accession numbers CP073033-40, NW_017264698-704 and NC_027860-66. Another genome of *O. polymorpha* DL-1 (GenBank: CP080316-22) was also used for syntenic comparison and SV detection, as its quality is higher than the reference genome of *O. polymorpha* DL-1 (RefSeq: NC_027860-66). Strand-specific RNA-seq data was (SRA: SRP124832) used to curate gene annotations of HU-11/CBS4732, NCYC495 and DL-1. The reads in this data correspond to the reverse complemented counterpart of transcripts.

## Supplementary information

Supplementary file 1: s1.txt

## Abbreviations

TR: tandem repeat
STR: short tandem repeat
LTR: long terminal repeat
mt: mitochondrial
nt: nucleotide
aa: amino acid
ORF: Open Reading Frame
CDS: Coding Sequence
SV: Structural Variation
(SNPs): single nucleotide polymorphisms
(InDels): insertions and deletions

## Declarations

### Ethics approval and consent to participate

Not applicable.

### Consent to publish

Not applicable.

### Availability of data and materials

The complete genome sequence of the Ogataea polymorpha strain CBS4732 ura3Δ (named HU-11) is available at the NCBI GenBank database under the accession numbers CP073033-40, in the project PRJNA687834.

### Authors’ contributions

SG conceived the project. SG and DW supervised the present study. JC assembled the HU-11/CBS4732 genome. JB and HF executed the experiments. SG, QS and TY analyzed the data. JC prepared the figures, tables and supplementary files. SG drafted the manuscript. SG, HW, WB and JR revised the manuscript. All authors have read and approved the manuscript.

## Acknowledgments

We appreciate the help equally from the people listed below. They are Wenjun Bu, Huaijun Xue, Dawei Huang, Yanqiang Liu, Bingjun He, Qiang Zhao, Zhen Ye and Xiufeng Jin from College of Life Sciences, Nankai University.

